# Predictability of evoked activity governs the geometry of spontaneous neural manifolds

**DOI:** 10.64898/2026.04.19.719506

**Authors:** Toshitake Asabuki, Claudia Clopath

## Abstract

Spontaneous neural activity reflects sensory experience yet occupies a lower-dimensional subspace than evoked responses. However, the principle governing which components of experience are incorporated into this intrinsic activity has remained unknown. Here, we propose that temporal predictability is one of the organizing principles that selects which evoked dimensions enter spontaneous neural manifolds. We show that local predictive synaptic plasticity provides a circuit mechanism that selectively embeds components of evoked activity that are predictable on the intrinsic timescale of recurrent dynamics, while excluding unpredictable fluctuations. As a consequence, the dimensionality of spontaneous activity is not fixed but depends on environmental timescales: rapidly fluctuating inputs are excluded, whereas slowly varying components are retained as contextual dimensions. This framework reproduces on- and off-manifold coding observed in visual cortex, and reconciles apparently conflicting developmental observations that spontaneous and evoked activity become both more similar and more geometrically distinct over maturation. Together, these results identify temporal predictability as a key principle linking environmental statistics to the geometry of intrinsic population dynamics.

## Introduction

Spontaneous neural activity, also known as ongoing activity, observed even in the absence of sensory input, is now recognized as a fundamental component of neural computation rather than mere background noise (Arieli et al., 1996; Tsodyks et al., 1999; Raichle, 2006; Fox &Raichle, 2007; Ringach, 2009; Uddin, 2020; Pezzulo et al., 2021). Across neural circuits, ongoing population activity exhibits highly structured spatiotemporal patterns that recapitulate key aspects of stimulus-evoked responses (Luczak et al., 2009; Luczak et al., 2013; Marguet & Harris, 2011). These findings suggest that neural circuits continuously generate internally organized activity that reflects learned models of the environment, even in the absence of sensory drive (Friston, 2010; Uddin, 2020; Pezzulo et al., 2021). Understanding how such intrinsic activity emerges from local circuit mechanisms is therefore crucial for explaining how the brain represents and predicts the world (Runfola et al., 2025).

A growing body of work suggests that spontaneous activity reflects internal models acquired through prior experience. In visual cortex, spontaneous activity occupies structured subspaces that overlap with those engaged during natural image processing (Kenet et al., 2003; Ringach, 2009). In the motor cortex, population activity is organized along low-dimensional manifolds that structure movement trajectories and constrain learning (Churchland et al., 2012; Shenoy et al., 2013; Gallego et al., 2017; Sadtler et al., 2014). In the hippocampus, place-cell ensembles re-express paths explored during behavior (Wilson & McNaughton, 1994; Foster & Wilson, 2006). Together, these studies suggest that recurrent circuits internalize structured regularities from sensory and behavioral experience through experience-dependent plasticity. Such internally organized dynamics may support replay and memory consolidation in some systems (Wilson & McNaughton, 1994; Foster & Wilson, 2006), and have also been interpreted as signatures of predictive coding or internal models of the environment (Rao & Ballard, 1999; Koren & Denève, 2017; Pezzulo et al., 2021; Dimakou et al., 2025).

Nevertheless, spontaneous patterns occupy a lower-dimensional subspace than evoked responses, even though both reflect the environment (Cunningham & Yu, 2014; Saxena & Cunningham, 2019; Stringer et al., 2019; Song et al., 2023; Iyer et al., 2022; Runfola et al., 2025). Intriguingly, several developmental studies present seemingly contradictory pictures of how spontaneous and evoked activity relate: spontaneous activity increasingly resembles evoked responses over postnatal development (Berkes et al., 2011), yet the two subspaces become more geometrically orthogonal with maturation (Avitan et al., 2021). This observation raises a deeper theoretical question: if spontaneous activity embodies an internal model of experience, what determines which components of experience are incorporated into spontaneous dynamics? Existing accounts based on metabolic efficiency or noise suppression (Sengupta et al., 2013) explain why dimensionality reduction may be advantageous, but do not specify the selection principle that determines which components of experience are incorporated and which are excluded.

Here, we propose that temporal predictability is a fundamental selection principle governing which components of evoked activity become incorporated into spontaneous neural manifolds. Prior work has shown that predictive learning can extract low-dimensional latent structure from the complex environment (Recanatesi et al., 2021; Levenstein et al., 2024; Asabuki & Fukai, 2020; Asabuki & Clopath, 2025). However, these frameworks did not address which subset of experience is selectively internalized into spontaneous activity—nor why spontaneous dimensionality is systematically lower than evoked dimensionality. We show that predictive plasticity selectively incorporates into spontaneous manifolds those components of input that are predictable on the intrinsic timescale of recurrent dynamics, while excluding fluctuations that cannot be recurrently reconstructed. This filtering principle explains why spontaneous and evoked activity can become more similar within the predictive subspace yet more geometrically separated in the full neural state space, thereby resolving the developmental paradox noted above. More broadly, our results reframe spontaneous population geometry as a direct readout of the brain’s implicit model of environmental timescales.

## Results

### Predictive learning selectively incorporates evoked dimensions into spontaneous manifolds

Spontaneous neural activity resembles stimulus-evoked responses yet occupies a lower-dimensional subspace, suggesting that it reflects only a subset of sensory experience. We hypothesized that this dimensional reduction arises from predictive learning, which selectively embeds predictable structure into recurrent dynamics while filtering out unpredictable fluctuations (Fig. 1A).

**Figure 1.**
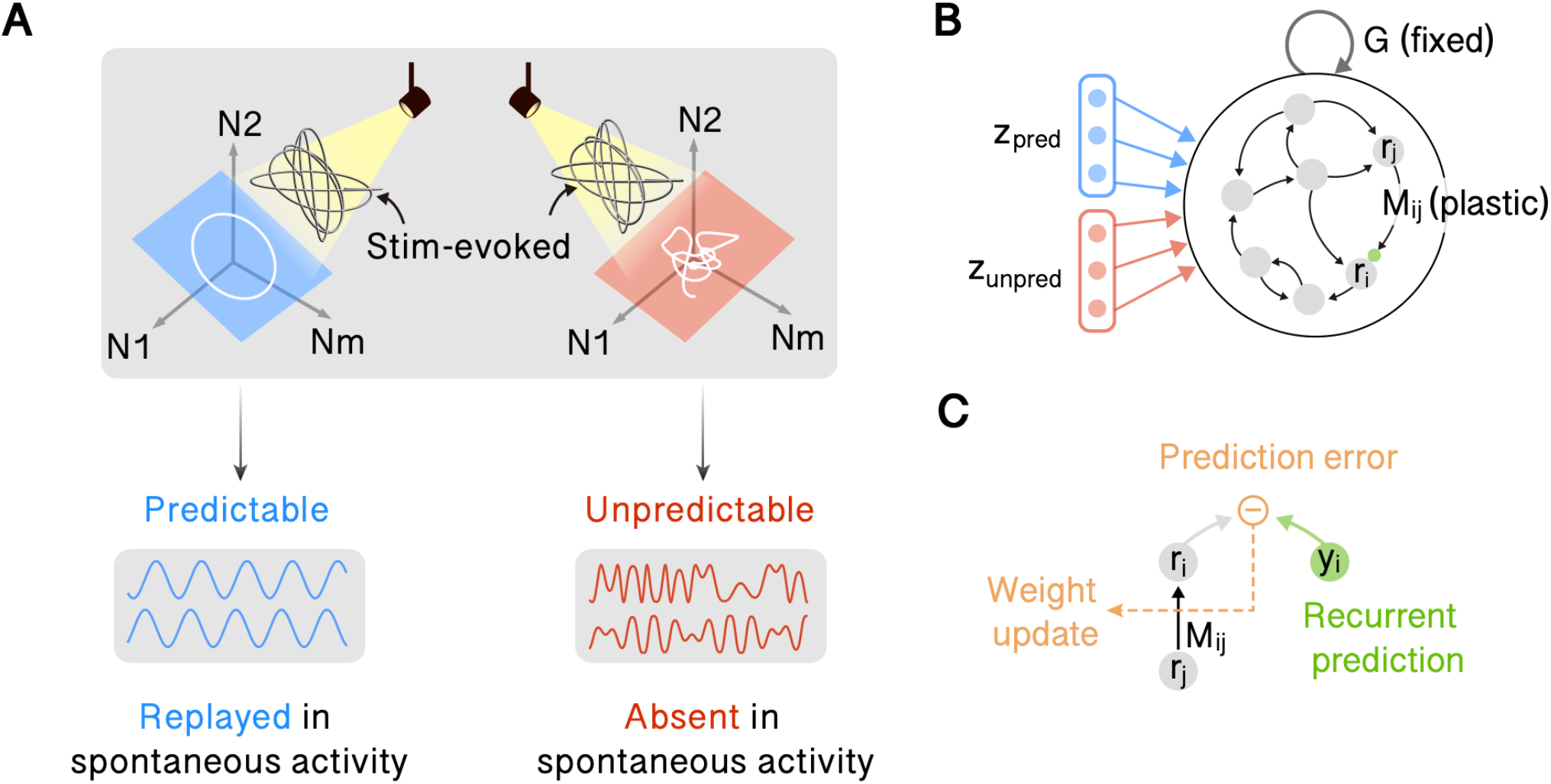
Predictive plasticity selectively embeds predictable dynamics into recurrent manifolds. **A** Conceptual illustration of predictable and unpredictable stimulus components. Sensory-evoked activity spans high-dimensional neural space (N1–Nm). Predictable components (blue) are embedded into a low-dimensional manifold that can be replayed during spontaneous activity. In contrast, unpredictable components (red) fail to stabilize within recurrent dynamics and are absent during spontaneous activity. **B** Network architecture. External inputs consist of predictable (z_pred, blue) and unpredictable (z_unpred, red) components. These inputs drive a recurrent network composed of a fixed connectivity G and plastic recurrent weights M. Only M is modified during learning, while G provides rich background dynamics. **C** Predictive learning rule. Each neuron forms a recurrent prediction (y_i_). The prediction error (r_i_ - y_i_) drives synaptic weight update according to the error-driven rule. This local plasticity rule embeds predictable temporal structure within recurrent connectivity while filtering out unpredictable fluctuations.

To test this hypothesis, we trained a rate-based recurrent neural network (RNN; Fig. 1B) using a local predictive plasticity rule (Asabuki & Clopath, 2025). The network consists of a fixed, strong random connectivity that generates high-dimensional chaotic dynamics, and a plastic recurrent component that is modified during learning. External inputs drive stimulus-evoked responses, and synaptic updates minimize the mismatch between these responses and recurrent predictions (Fig. 1C).

More precisely, synaptic changes are driven by the product of a prediction-error term (the difference between a neuron’s current activity and its recurrent prediction) and a presynaptic activity term. Importantly, both signals are locally available at each synapse: the prediction error is computed at the postsynaptic neuron based on its own activity and the recurrent inputs it receives, while the presynaptic activity reflects the input from the corresponding afferent neuron. Both signals undergo temporal filtering prior to influencing plasticity, analogous to postsynaptic and presynaptic integration in biological neurons. The recurrent prediction integrates recent recurrent input over a short period, and the presynaptic activity is converted into an eligibility trace. This learning rule is fully local, consistent with biological constraints on synaptic plasticity.

We first considered a three-dimensional input signal. Two dimensions, *z*_1_ and *z*_2_, traced a structured trajectory representing canonical predictable dynamics, whereas the third dimension, *z*_3_, followed an irregular, unpredictable time course (Fig. 2A, top). During training, the prediction error decreased monotonically (Supplementary Fig. 1), indicating convergence. After learning, we removed the external drive and examined spontaneous activity. Structured spontaneous activity persisted following removal of the external inputs and remained highly similar to the pattern observed during evoked conditions (Fig. 2A, bottom; Fig. 2B). The network spontaneously generated a trajectory confined almost entirely to the two-dimensional subspace defined by the predictable inputs, whereas activity along the unpredictable dimension decayed to baseline (Fig. 2C).

**Figure 2.**
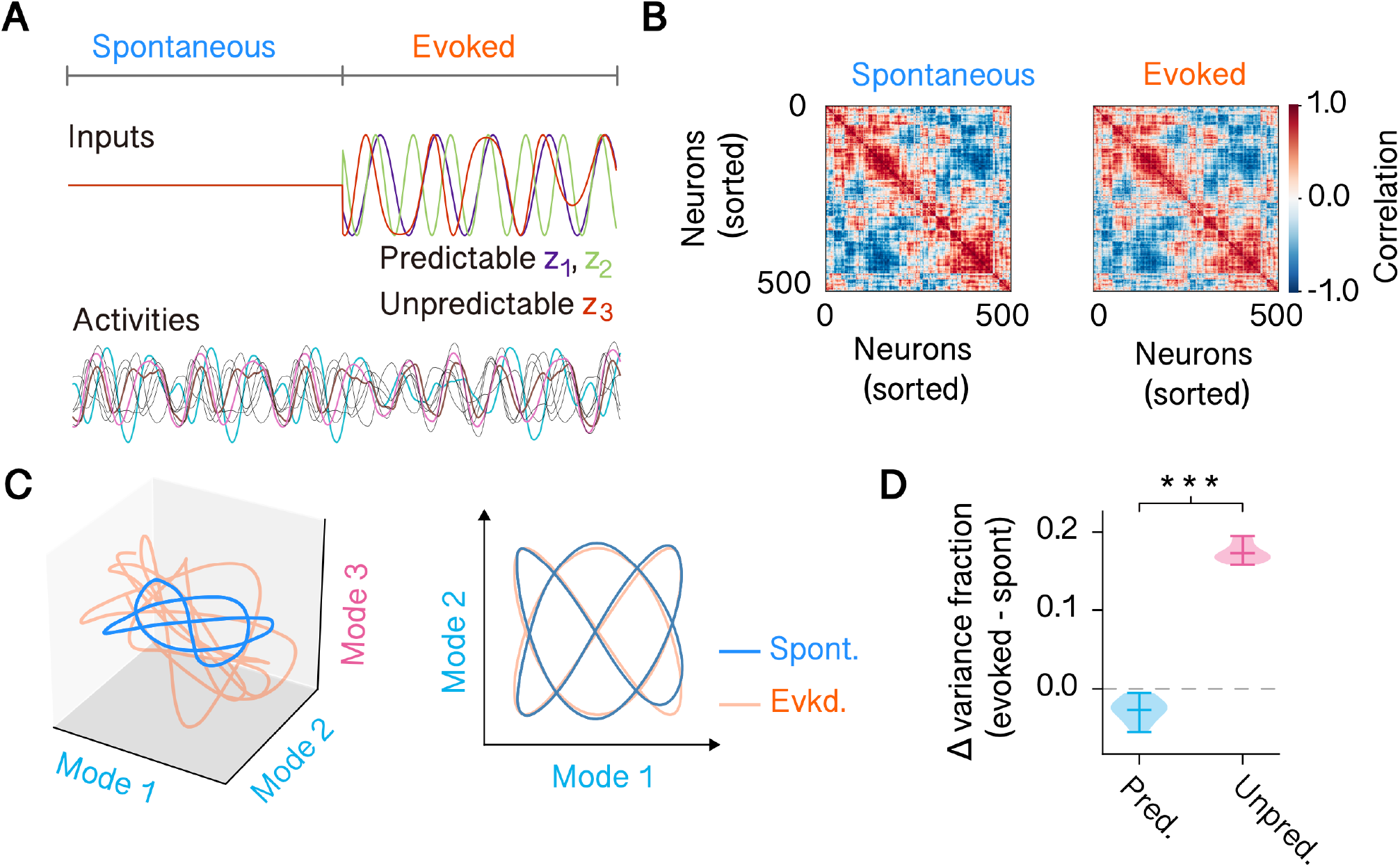
Predictive plasticity selectively stabilizes predictable population modes. **A** Example input signals and network activity after learning. During the spontaneous phase (left), no external inputs are present and the network evolves autonomously. During the evoked phase (right), temporally structured inputs are given to the network. Top: example input signals. Bottom: representative neural activities from multiple neurons after learning. **B** Pairwise correlation structure during spontaneous and evoked activity. Neurons are sorted according to spontaneous activity correlations. The block structure observed during spontaneous activity is partially preserved during evoked responses, indicating stabilization of intrinsic population modes. **C** Low-dimensional population dynamics. Left: trajectories projected onto the three neural modes during spontaneous (blue) and evoked (orange) activity. Right: projection onto the first two neural modes. Within this subspace, spontaneous and evoked trajectories exhibit similar dynamical structure, indicating that predictable components are embedded within intrinsic population modes. **D** Change in variance fraction between evoked and spontaneous activity for predictable (Pred.) and unpredictable (Unpred.) input components across 10 independent simulations. Predictable dimensions show minimal variance change, whereas unpredictable components exhibit significant expansion beyond the spontaneous manifold. P values were calculated using a two-sided Welch’s t test (^***^P < 0.001).

To quantify how stimulus presentation reshaped the geometry of population activity, we compared the fraction of the variance along predictable and unpredictable input dimensions during evoked and spontaneous activity. Across 10 independent simulations, predictable dimensions exhibited minimal change in explained variance between spontaneous and evoked states, indicating that these components were already embedded within the spontaneous manifold (Fig. 2D). In contrast, unpredictable dimensions showed a significant increase in variance during evoked activity, reflecting a transient expansion beyond the spontaneous subspace.

Next, we tested how temporal filtering in the plasticity rule contributes to the formation of spontaneous dynamics that reflects the predictable part of evoked activity. To this end, we removed each low-pass component in turn and examined the resulting network behavior. When postsynaptic integration was removed, recurrent prediction followed current activity too quickly, preventing the network from capturing slowly varying temporal regularities in the input. The resulting spontaneous activity became high-dimensional and irregular, with no trace of previously experienced trajectories (Supplementary Fig. 2A). A similar disruption occurred when the presynaptic low-pass filter was removed. Because synaptic updates responded to instantaneous fluctuations rather than stable correlations, the network learned a distorted internal model. Although spontaneous activity retained low-dimensional structure, it failed to align with the evoked manifold (Supplementary Fig. 2B). Only when both filters were present did the network successfully internalize predictable temporal structure and autonomously reproduce it after training.

Together, these results demonstrate that predictive learning embeds externally driven dynamics within spontaneous activity confined to a low-dimensional manifold that captures the predictable aspects of sensory experience while discarding unpredictable signals.

### Temporal predictability determines the dimensionality of spontaneous manifolds

The above results suggest that the network learns to distinguish predictable signals from unpredictable noise. However, when apparently irregular signals contain slow temporal correlations, they may still influence the structure of the learned spontaneous activity. We therefore hypothesized that the timescale of input fluctuations determines whether a signal component is incorporated into or excluded from the network’s spontaneous repertoire.

To test this hypothesis, we used the same three-dimensional input structure as before. We varied the timescale of fluctuations in the irregular component, z_3_(t), while keeping its overall statistics and amplitude constant. To simplify the analysis, we replaced the predictable component (z_1_, z_2_) with a circular trajectory rather than the more complex structured pattern used previously. The unpredictable component, z_3_(t), was generated with two frequencies, one fast (Fig. 3A) and one slow (Fig. 3B). The high-frequency condition corresponded to the setting used in the previous experiment.

**Figure 3.**
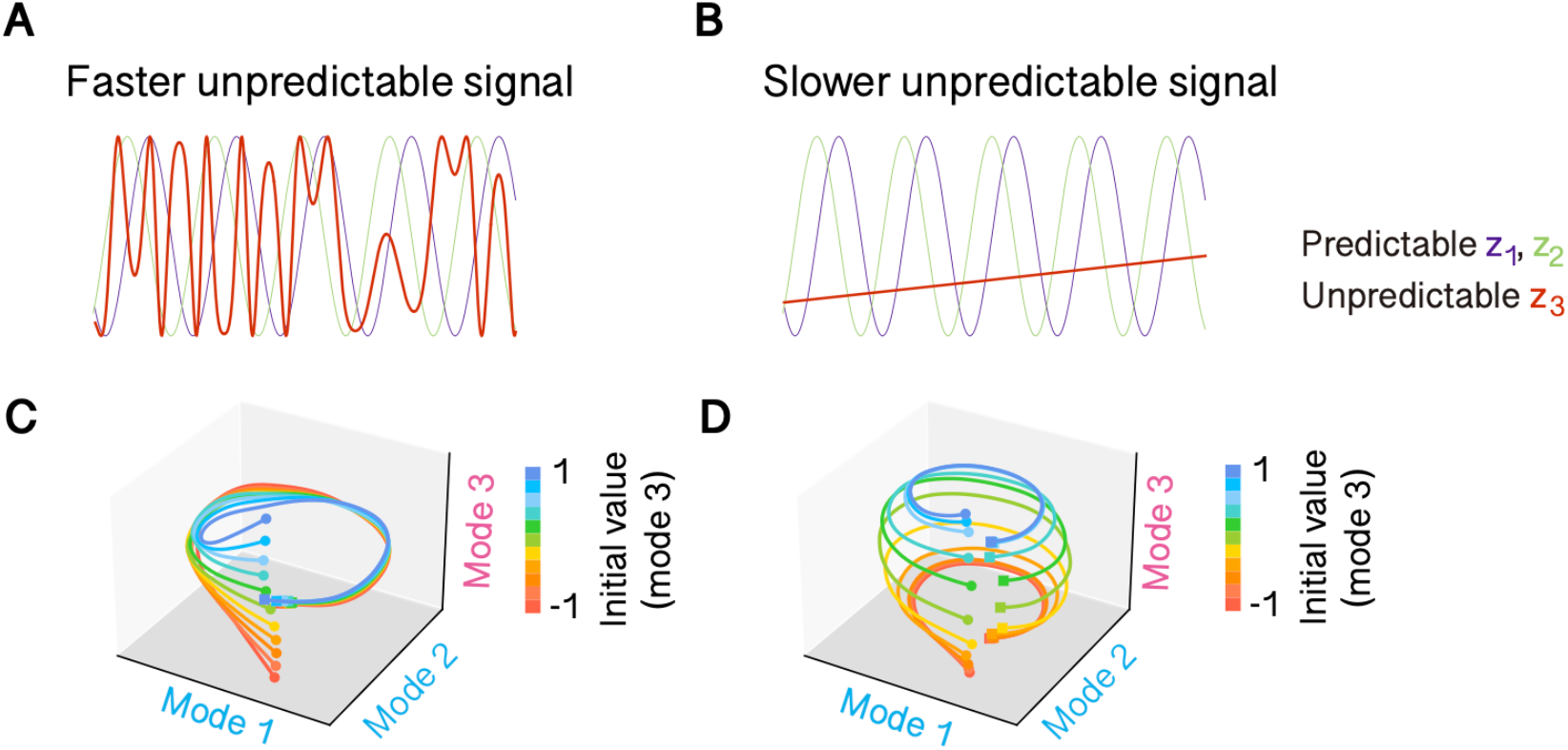
Embedding of unpredictable signals depends on their temporal timescale. **A-B** Example input signals in which the unpredictable component varies on a slower (A) or faster (B) timescale relative to the predictable component. Colored traces illustrate different initial values along the third population mode. **C-D** Low-dimensional population trajectories projected onto the first three principal modes after learning. Colors indicate different initial values along mode 3. When the unpredictable signal varies rapidly (C), it becomes excluded from the low-dimensional spontaneous population structure. In contrast, when the unpredictable signal varies slowly (D), it becomes embedded within intrinsic dynamics and induces expansion along additional dimensions.

After training the network under each condition, we removed the external drive and initialized it from multiple initial states. When trained with high-frequency fluctuations, the network’s spontaneous activity always converged on the same two-dimensional manifold defined by the predictable circular drive (Fig. 3C). In contrast, when the input fluctuations were slow, the learned spontaneous activity retained the initial offset along the z_3_ axis while still following the circular trajectory in (z_1_, z_2_) (Fig. 3D). Thus, different initial conditions led to parallel circular trajectories displaced along z_3_, forming a three-dimensional cylindrical manifold.

Taken together, these results demonstrate that the dimensionality of spontaneous dynamics is not fixed but rather adapts to the timescale of external inputs. The network selectively internalizes predictable signal components on its intrinsic timescale, preserving slowly varying inputs as stable contextual dimensions.

### Geometric separation enables multiple predictable manifolds to coexist with minimal interference

Having shown that predictive plasticity selects which evoked dimensions are incorporated into a spontaneous manifold, we next asked whether the same principle can organize multiple predictable structures within a single recurrent circuit. If predictability indeed organizes spontaneous population geometry, then distinct predictable structures should be stored as separable manifolds whose retrieval depends on their geometric separation. To test this idea, we trained the network on two distinct trajectories presented in alternating blocks. One input traced a circular trajectory confined to a specific two-dimensional subspace (z_1_, z_2_), and the other followed a different trajectory confined to a separate, orthogonal subspace (z_4_, z_5_) (Fig. 4A). Each trajectory was combined with an unpredictable component along z_3_ or z_6_, respectively. During training, these two input patterns alternated across epochs (Fig. 4B). By embedding the dynamics in orthogonal subspaces, we hypothesized that the learning rule would modify largely independent sets of synaptic weights, thereby minimizing interference between the two learned internal models.

**Figure 4.**
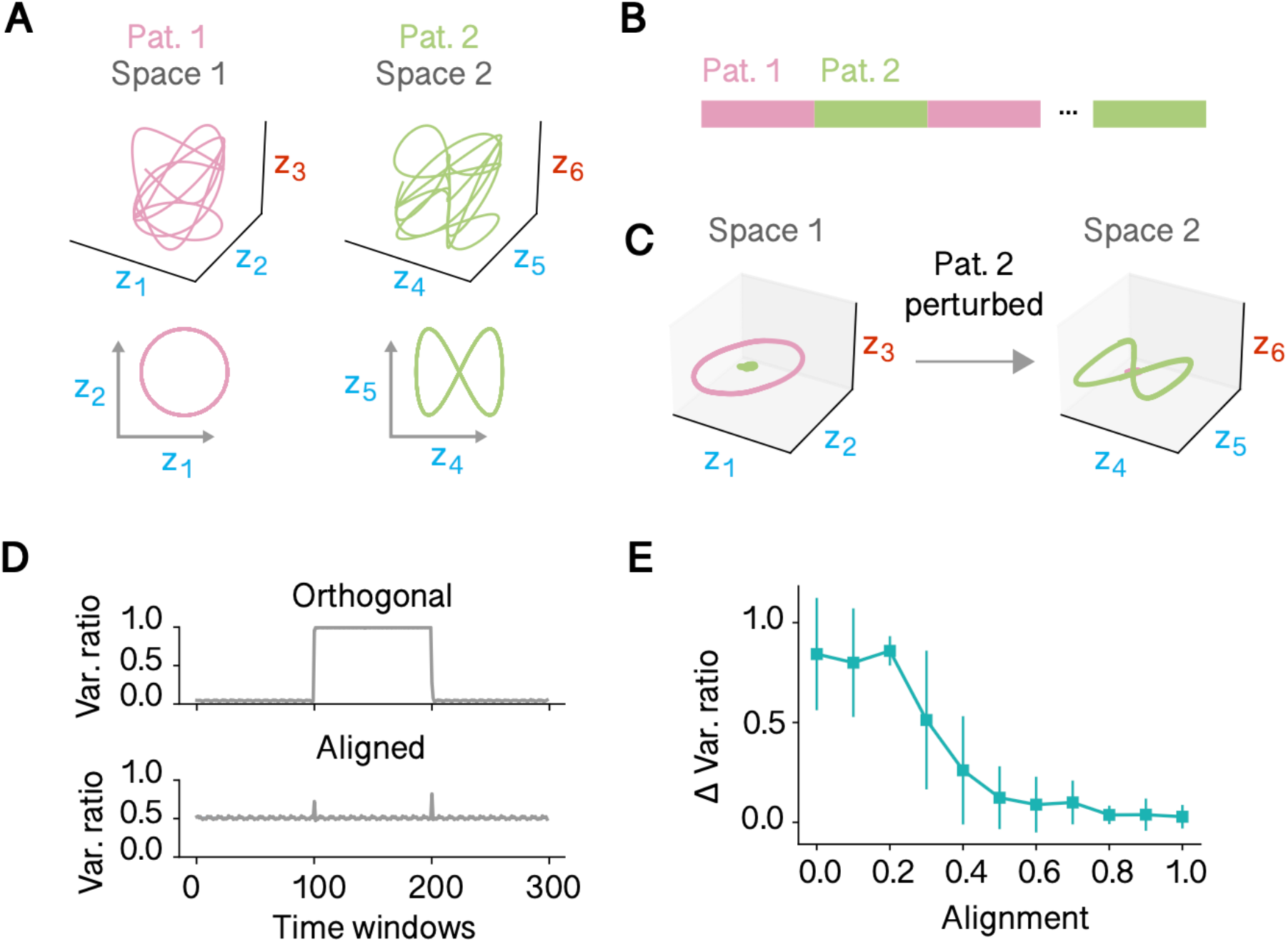
Predictive plasticity organizes multiple spontaneous manifolds through geometric separation. **A** Low-dimensional population trajectories for two learned patterns (Pattern 1 and Pattern 2). Each pattern contains both predictable and unpredictable components. After learning, the predictable component of each pattern is embedded within a distinct population subspace (Space 1 and Space 2). **B** Alternating presentation of Pattern 1 and Pattern 2 during training. **C** Pattern switching via subspace-specific perturbation. During spontaneous activity, the network autonomously replays the predictable component of one learned pattern (e.g., Pattern 1). A brief pulse perturbation delivered along the predictable axis of Pattern 2 induces a transition of population activity into the manifold corresponding to Pattern 2, where its predictable component is replayed. **D** Dynamics of variance ratio (VR) quantifying subspace dominance during spontaneous switching. When the two predictable manifolds are orthogonal, VR alternates between values near 0 and 1, indicating that population activity switches cleanly between the two manifolds. In contrast, when the manifolds are aligned, VR remains near 0.5, reflecting mixed activity and reduced subspace separation. **E** Dependence of variance ratio separation on subspace alignment. Alignment between the two predictable manifolds was systematically varied. Error bars indicate mean ± s.e.m. across 10 independent simulations. Increasing alignment reduces subspace separation and increases interference.

After training, we tested whether the network’s spontaneous activity could distinctly represent these two trajectories. To this end, we briefly perturbed the trained network with a transient pulse along one of the predictable dimensions. Strikingly, the network could rapidly and reliably switch between the two internally generated dynamical states based solely on such a transient cue (Fig. 4C). In the spontaneous regime, presenting the pulse associated with the circular trajectory initiated self-sustained replay of the corresponding dynamics with minimal activation in the orthogonal subspace representing the other trajectory (see Fig. 4C, left), and vice versa (see Fig. 4C, right).

To examine how the separability of learned subspaces influences memory interference further, we systematically varied the degree of overlap between the two trajectory subspaces. For convenience, we refer to the subspaces corresponding to (z_1_, z_2_) and (z_4_, z_5_) as space 1 and space 2, respectively. We gradually increased the overlap between the two subspaces by blending space 2 toward space 1 while keeping the additional dimensions (z_3_, z_6_) orthogonal to both. This manipulation allowed us to control the amount of shared structure between the two learned manifolds continuously.

After training under each overlap condition, we evaluated the network’s performance to switch between the two spontaneous trajectories. When the two subspaces were fully orthogonal, the network robustly maintained and switched between both manifolds (Fig.4D, top). In contrast, when the subspaces completely overlapped, spontaneous switching was no longer possible; the network’s activity converged to a single shared attractor (Fig.4D, bottom). As the overlap increased, switching performance remained stable for small overlaps but gradually declined beyond a certain range (Fig. 4E). Note that, performance declined steeply beyond moderate overlap, suggesting limited tolerance for representational interference between stored manifolds.

Together, these results show that predictive plasticity can organize multiple internal models as distinct spontaneous manifolds within a single recurrent network. This capacity depends critically on the geometric separability of their predictable subspaces, revealing how population geometry controls both retrieval and interference between learned dynamical states.

### Predictive manifold selection explains on- and off-manifold coding in cortex

So far, we have shown that predictive plasticity selectively embeds temporally predictable dynamics into low-dimensional spontaneous manifolds. We next asked whether this principle of predictive manifold selection can account for the experimentally observed dissociation between on- and off-manifold coding in visual cortex, where spontaneous population activity during slow-wave sleep (SWS) defines an intrinsic manifold, whereas awake activity recruits within-manifold dimensions for behavior-related signals and orthogonal dimensions for stimulus-related signals (Fig. 5A; de Oliveira et al., 2022).

**Figure 5.**
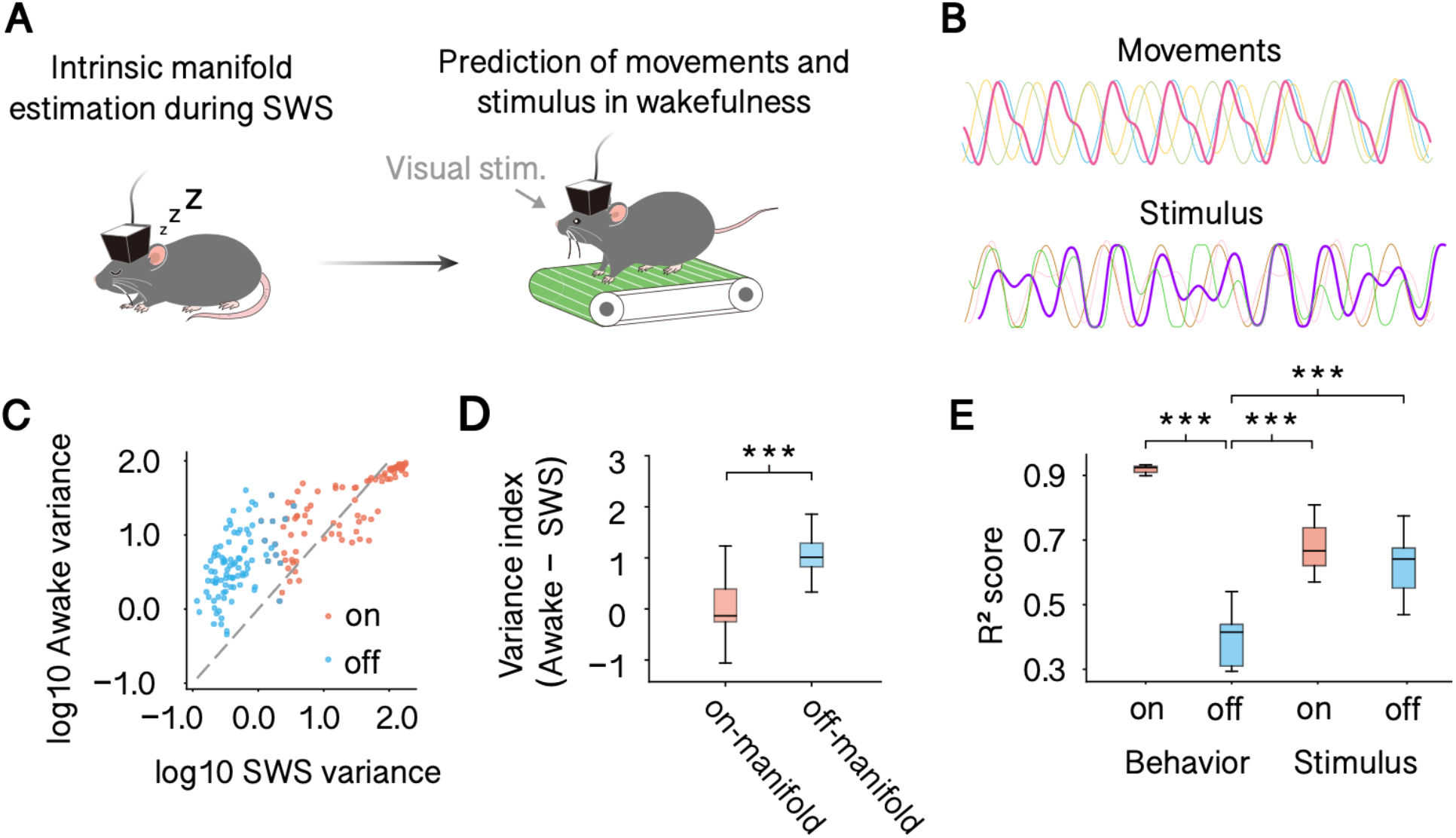
Predictive plasticity reproduces on- and off-manifold coding signatures. **A** Experimental setting of de Oliveira et al., 2022. During slow-wave sleep (SWS), spontaneous activity defines an intrinsic low-dimensional manifold. During wakefulness, behavior- and stimulus-related activity are expressed relative to this intrinsic manifold. **B** Modeled behavioral and stimulus signals used as inputs. Movements consist of predictable components. Stimulus signals contain both predictable and unpredictable components. **C** Comparison of variance between spontaneous (SWS-like) and evoked (awake) activity for individual population modes. Each point represents a mode. Predictable (on-manifold) modes show similar variance during spontaneous and evoked activity, whereas additional variance emerges in off-manifold dimensions during evoked responses. **D** Variance index (Awake - SWS) for on-manifold and off-manifold dimensions. On-manifold dimensions exhibit minimal variance change, whereas off-manifold dimensions show increased variance during evoked activity. **E** Decoding performance (R^2^ score) from on- and off-manifold projections for behavioral and stimulus signals. Behavioral signals are predominantly decodable from on-manifold dimensions, whereas stimulus signals engage both on- and off-manifold subspaces. In D and E, data are shown as mean ± s.e.m.; _***_P < 0.001 over 20 independent simulations.

To test whether our model can explain this experimental finding, we trained the network on a higher-dimensional input structure. Behavioral signals consisted of four temporally predictable components, whereas stimulus signals contained two predictable and two unpredictable components (Fig. 5B). This design allowed us to assess whether the learned spontaneous manifold selectively captures predictable structure while evoked responses recruit additional dimensions to represent unpredictable stimulus variability.

After training, we extracted the spontaneous manifold from autonomous network activity and defined the on-manifold subspace using its leading principal components. For comparison, an equal number of low-variance components defined the off-manifold subspace (Methods). We then examined how variance along individual population modes changed between spontaneous (SWS-like) and evoked (awake-like) activity (Fig. 5C). Predictable, on-manifold modes exhibited comparable variance across states, indicating that these dimensions were already embedded within the spontaneous dynamics. In contrast, additional variance emerged selectively in off-manifold modes during evoked activity, reflecting a state-dependent expansion beyond the intrinsic manifold. To quantify this effect, we computed a variance index (Awake - SWS) separately for on- and off-manifold dimensions (Fig. 5D). On-manifold dimensions showed minimal variance change, whereas off-manifold dimensions displayed a significant increase during evoked responses. Thus, evoked activity showed a structured expansion: variance was largely preserved within the spontaneous manifold while increasing preferentially in off-manifold dimensions.

To test the functional content carried by each subspace, we next assessed decoding performance using linear readouts (Fig. 5E). Behavioral signals were robustly decodable from the on-manifold projection but not from the off-manifold projection, indicating that predictable self-generated dynamics are embedded within the spontaneous manifold. In contrast, stimulus signals required both subspaces for accurate reconstruction: decoding performance was highest when including off-manifold dimensions, consistent with the recruitment of orthogonal directions by unpredictable stimulus components.

Together, these results demonstrate that predictive plasticity reproduces the experimentally observed dissociation between on- and off-manifold coding. Predictable behavioral structure remains confined within the spontaneous manifold, whereas unpredictable stimulus variability selectively engages additional dimensions beyond it.

## Discussion

Spontaneous neural activity is not a passive echo of prior sensory input, but a geometrically organized internal model of the subset of experience that recurrent circuits can autonomously sustain. Here we show that predictive synaptic plasticity provides a circuit-level principle that sculpts this geometry. By selectively stabilizing temporally predictable components of sensory input, recurrent networks compress high-dimensional experience into low-dimensional manifolds that can be autonomously sustained. Rather than minimizing dimensionality for efficiency alone, predictive learning allocates population activity into a self-consistent predictive subspace while excluding unpredictable fluctuations. Importantly, the exclusion of unpredictable components is not a design choice but an inevitable consequence of predictive learning operating on finite timescales. Thus, population geometry reflects not the full structure of sensory inputs, but the subset that can be internally predicted and sustained by recurrent dynamics.

This dependence arises because spontaneous activity can internalize only those components of past input that are predictable on the intrinsic timescale of recurrent dynamics. The dimensionality of spontaneous manifolds is therefore not fixed, but selected by the temporal statistics of environmental variability. Rapid fluctuations are excluded because they cannot be recurrently reconstructed, whereas slowly varying inputs are retained as contextual dimensions. Spontaneous manifold geometry thus reflects not only the complexity of past inputs but also the temporal structure of uncertainty present during learning, consistent with theoretical views linking neural dynamics to prediction and environmental statistics (Huang & Rao, 2011; Chalk et al., 2018; Koren & Denève, 2017). In this sense, population geometry encodes predictability structure rather than stimulus identity per se.

Our framework makes experimentally tractable predictions about how environmental statistics shape spontaneous population geometry. If spontaneous manifold dimensionality depends on the temporal structure of sensory noise encountered during learning, then manipulating environmental noise correlations should systematically reshape spontaneous activity. This prediction resonates with prior views that spontaneous activity reflects learned environmental statistics. Animals exposed to slowly varying structured noise should exhibit higher-dimensional spontaneous manifolds than those trained under rapidly fluctuating noise conditions (Berkes et al., 2011; Orbán et al., 2016; Fiser et al., 2010; Recanatesi et al., 2021). Such experiments would provide a direct test of whether temporal predictability, rather than coding efficiency alone, determines which evoked dimensions are incorporated into spontaneous manifolds. Likewise, altering sensory experience should reshape the balance between on- and off-manifold representations observed during evoked activity, by changing the relationship between predictable and unpredictable stimulus components (Fiser et al., 2004; Stringer et al., 2019). Together, these experiments would directly test whether spontaneous geometry tracks environmental predictability.

Our framework reconciles seemingly contradictory developmental observations. Berkes et al. (2011) reported that spontaneous activity increasingly resembles evoked responses over development, whereas Avitan et al. (2021) found that the two become more orthogonal. We propose that predictive plasticity selectively embeds temporally predictable components of sensory experience into recurrent connectivity. Within the low-dimensional predictive subspace defined by these stable regularities, spontaneous activity converges toward evoked structure. At the same time, unpredictable and fast-varying components are excluded from recurrent dynamics, producing geometric divergence in the full neural state space. Cortical development therefore might reflect geometric refinement of predictive structure, with predictable components stabilized and unpredictable variability progressively excluded.

Our framework also provides a geometric account of on- and off-manifold coding reported in cortex. Because spontaneous activity reflects a stabilized predictive subspace, behavior-related signals that can be internally generated remain largely confined within this manifold. In contrast, stimulus-driven variability that cannot be reconstructed from recurrent dynamics necessarily recruits additional dimensions. This separation arises as a consequence of predictive learning rather than from an explicit routing mechanism, offering a complementary perspective to hierarchical predictive coding theories that emphasize error propagation (Friston, 2010; Huang & Rao, 2011; Bastos et al., 2012), as well as to population geometry studies demonstrating orthogonal subspaces for distinct computations (Elsayed et al., 2016).

At the physiological level, the proposed mechanism is consistent with known properties of cortical microcircuits. The dual temporal filtering of pre- and postsynaptic signals resembles dendritic and synaptic integration processes and may arise from biophysical mechanisms such as NMDA receptor-dependent currents, membrane time constants, and local inhibitory control of recurrent excitation. In this view, spontaneous manifolds arise through continuous local prediction within recurrent networks, without requiring replay or global coordination. Such a mechanism may contribute to the emergence of intrinsic timescales, metastable state transitions, and context-dependent reactivation observed in cortical dynamics (Deco et al., 2013; Rabinovich et al., 2008; Litwin-Kumar & Doiron, 2012; Breakspear, 2017; Mazzucato et al., 2019).

Together, these findings suggest that spontaneous neural activity represents the self-consistent core of prior experience: the subset of environmental dynamics that a circuit can internally sustain. Predictive plasticity may therefore provide a general circuit principle by which environmental statistics determine which evoked dimensions are consolidated into spontaneous population geometry.

## Methods

### Neural network model

Our network consists of *N* rate-based neurons, mutually connected with the recurrent connections. Here, we considered two types of recurrent connections: strong and fixed connectivity G and initially weak yet plastic connectivity M (Asabuki and Clopath, 2025). The strength of each connection was generated by a Gaussian distribution, and unless other specified, with zero-mean and the standard deviation of 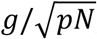 with (*p, g*) *=* (0.1, 1.2) for G and (*p, g*) = (1.0, 0) for initial strength of M. Here, *g* is a gain of connection, leading chaotic spontaneous activity with *g >* 1 (Sompolinsky et al., 1988), thus the fixed recurrent connectivity G generates initial chaotic spontaneous activity. The dynamics of membrane potential with the external input ***I*** was governed by the following equation:

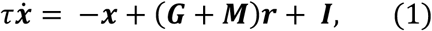

where ***x*** is a membrane potential and ***r*** is a firing rate of neuron, defined as:

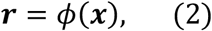

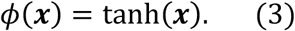

The parameter *τ* is a membrane time constant, which we set as *τ* = 10 (ms).

### Input signals

The external input ***I***(*t*) ∈ ℝ^*N*^ was defined as a linear projection of a low-dimensional signal vector *z*(*t*) *=* [z_1_(*t*), z_2_(*t*), *z*_3_(*t*)]^T^ onto the neural population through a fixed input matrix *W*_*z*_ ∈ ℝ ^N×3^ :

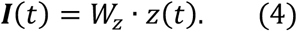

Each column of *W*_*z*_ represented an orthogonal input channel. In all figures except Figure 4, we constructed the input weight matrix *W*_*z*_ as follows. We first generated a random matrix *R* ∈ ℝ^N×^_3_ with uniformly distributed elements in [-0.5, 0.5] and performed QR decomposition, obtaining an orthogonal basis *Q* (*Q*^*T*^*Q = I*_3_). We then defined the input weight matrix as

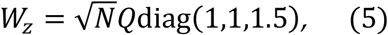

where diag is a diagonal matrix. The three components of *Z*(*t*) corresponded to distinct temporal features of the environment: z_1_(*t*) and z_2_(*t*) represented predictable, structured dynamics, whereas z_3_(*t*) carried irregular, unpredictable fluctuations.

For the simulations shown in Fig. 2, the predictable components traced a structured yet nontrivial trajectory in the z_1_−z_2_ plane:

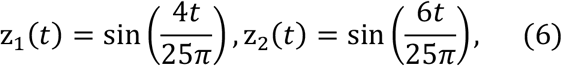

while the unpredictable component z_3_(*t*) followed a slowly varying and irregular function,

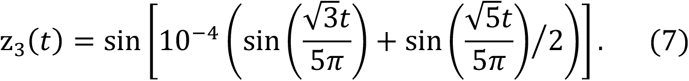

This combination yielded a complex input trajectory containing both temporally regular and irregular components, allowing the network to learn the predictable portion of the dynamics while ignoring the unpredictable component.

For Figs. 3–5, we used a simplified input structure in which the predictable components formed a circular trajectory:

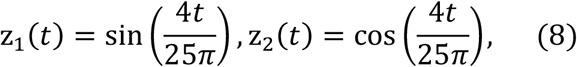

and the same irregular signal was used for z_3_(*t*) as above.

### The learning rule

The synaptic plasticity rule was designed to predict the externally driven network dynamics. To this end, we defined the recurrent prediction ***y*** evolve as a low-pass filtered version of recurrent synaptic current ***Mr***:

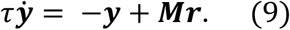

The learning rule aimed to minimize the mismatch between the recurrent prediction and the actual neural activity:

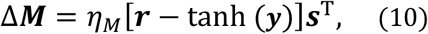

where ***s*** denotes the low-pass filtered presynaptic activity:

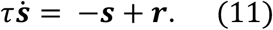

Here, *η*_*M*_ is the learning rate. This rule drives ***M*** to generate recurrent predictions that match the input-driven responses, thereby filtering out the unpredictable components from the external drive. After training, only the internally generated dynamics that capture the predictable structure of the inputs persist in the spontaneous activity.

### Analysis of effect of timescales

To examine how the timescale of input fluctuations influences the formation of spontaneous manifolds (Fig. 3), we reduced the frequency of z_3_(*t*):

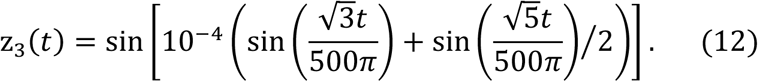

This modification preserved the amplitude of the signal while producing slow, quasi-stationary fluctuations. After training, we tested how the network’s spontaneous dynamics depended on the initial value of z_3_. We initialized the network from ten distinct starting conditions, where z_3_took equally spaced values between -1 and +1 while z_1_ and z_2_ were set to 1 and 0, respectively.

### Learning multiple subspaces

The network received six external input channels: three for manifold 1 (z_1_(*t*), z_2_(*t*), *Z*_3_(*t*)) and three for manifold 2 (z_4_(*t*), z_5_(*t*), *Z*_6_(*t*)). Each channel projected onto the network via an input vector *w*_*i*_ ∈ ℝ^*N*^(*i* = 1, …,6)). We generated six random vectors, orthogonalized them, and used the first three (w_1_, w_2_, *w*_3_) as the basis of manifold 1. The remaining three (w_4_, w_5_, *w*_6_) were constructed to share a controlled degree of overlap with manifold 1, defined by a mixing coefficient *α* (0 = orthogonal, 1 = identical):

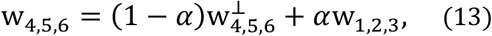

where 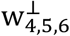 denote the orthogonalized components relative to w_1,2,3_. All weight vectors were normalized to have 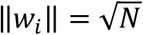.

During learning, the two input sets alternated every 10 s. For manifold 1, the external signals were Eq.(8) and Eq.(6) for manifold 2. During testing, the network initially evolved under spontaneous conditions. We then applied a brief 50-ms pulse to one of the input subspaces to cue the corresponding manifold. After cue removal, all inputs were set to zero.

We quantified subspace dominance by computing a variance ratio within 100 ms sliding windows of the network activity r_*t*_, capturing how the relative variance across the two manifolds evolved over time:

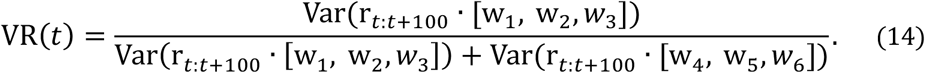

An increase in the variance ratio indicated that network activity was confined within manifold 1, whereas a decrease reflected a transition toward manifold 2.

### Analysis of on- and off-manifold coding

We quantified how the network represented predictable and unpredictable information across on- and off-manifold subspaces. The input signals consisted of three components: z_1_ (predictable behavioral dynamics), z_2_ (predictable sensory component of visual stimuli), and z_3_ (unpredictable sensory fluctuations).

After training, we simulated the network in two conditions—spontaneous (SWS-like) with all inputs set to zero, and evoked (awake-like) with all inputs active. We defined the spontaneous manifold by applying principal component analysis (PCA) to spontaneous population activity. The top principal components explaining 99% of cumulative variance were defined as the on-manifold subspace, and an equal number of components from the tail of the variance spectrum were defined as the off-manifold subspace, matched in dimensionality.

We projected evoked activity onto these two subspaces and used ridge regression (regularization coefficient = 10) to decode each input signal from the neural population activity. Decoding accuracy was quantified by the coefficient of determination (*R*^2^) for (i) the behavioral signal (z_1_) and (ii) the sensory signal, computed as the average decoding accuracy for z_2_ and z_3_. We compared decoding performance between on- and off-manifold subspaces using Welch’s two-tailed t-tests.

To visualize variance changes between spontaneous and evoked states, we computed the log-transformed variance of each PCA dimension and defined a variance index as

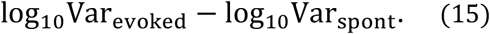

We compared the distributions of variance indices between on- and off-manifold dimensions. We also performed Welch’s two-tailed t-tests to assess group-level differences in decoding accuracy and variance indices.

### Simulation details

All simulations were performed in customized Python3 code written by TA with numpy 2.3.5 and scipy 1.17.0. Differential equations were numerically integrated using an Euler method with integration time steps of 1 ms.

## Acknowledgments

The authors thank Juan Alvaro Gallego for their valuable comments on our manuscript. This work was supported by Wellcome Trust 200790/Z/16/Z, Simons Foundation 564408, EPSRC EP/R035806/1, ERC MotorAdapt 101169605 to C.C.

## Author Contributions

T.A. and C.C. conceived the study and wrote the paper. T.A. performed the simulations and data analyses.

## Supplementary Figures

**Supplementary Figure 1:**
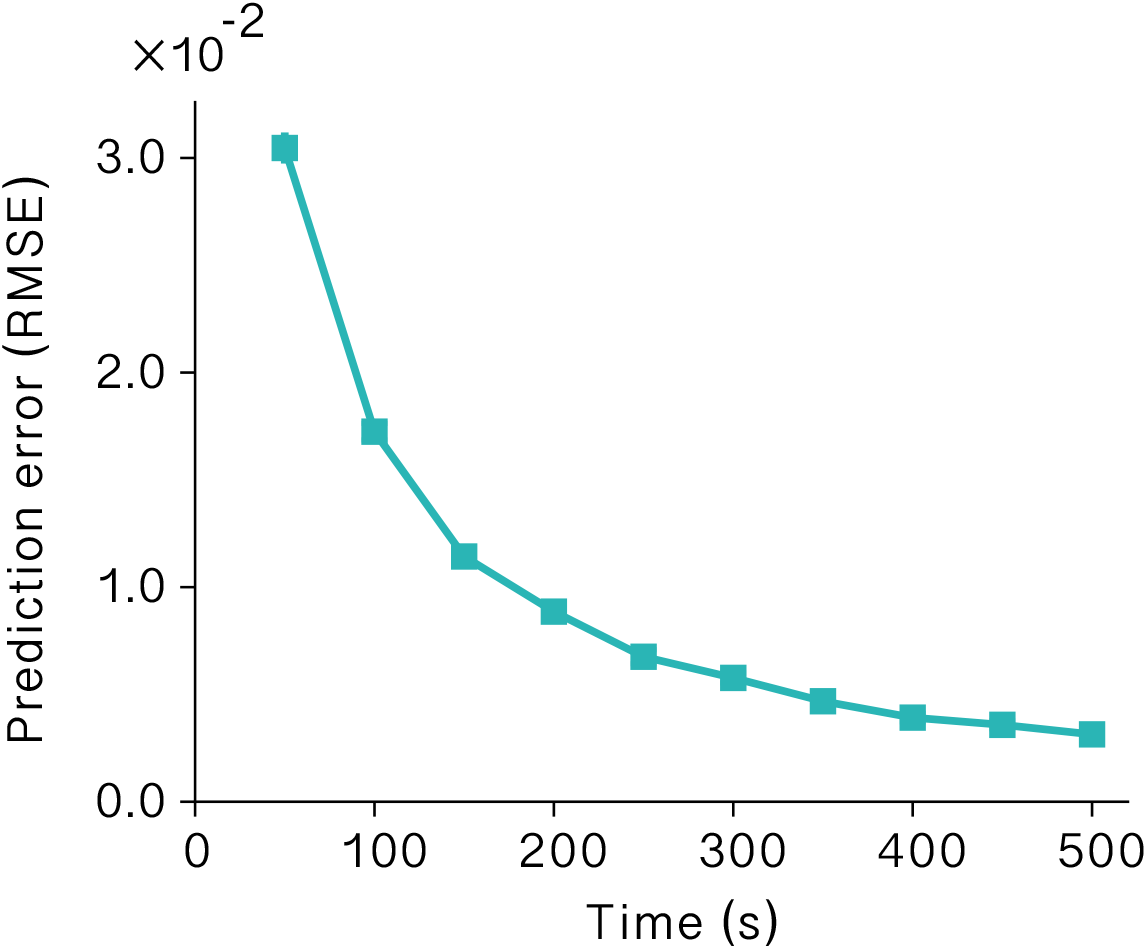
Model performance during learning. Performance of our model during learning, quantified as the root mean squared error, is shown by the dashed line. Error bars (invisible) stand for s.d.s over 20 independent simulations.

**Supplementary Figure 2:**
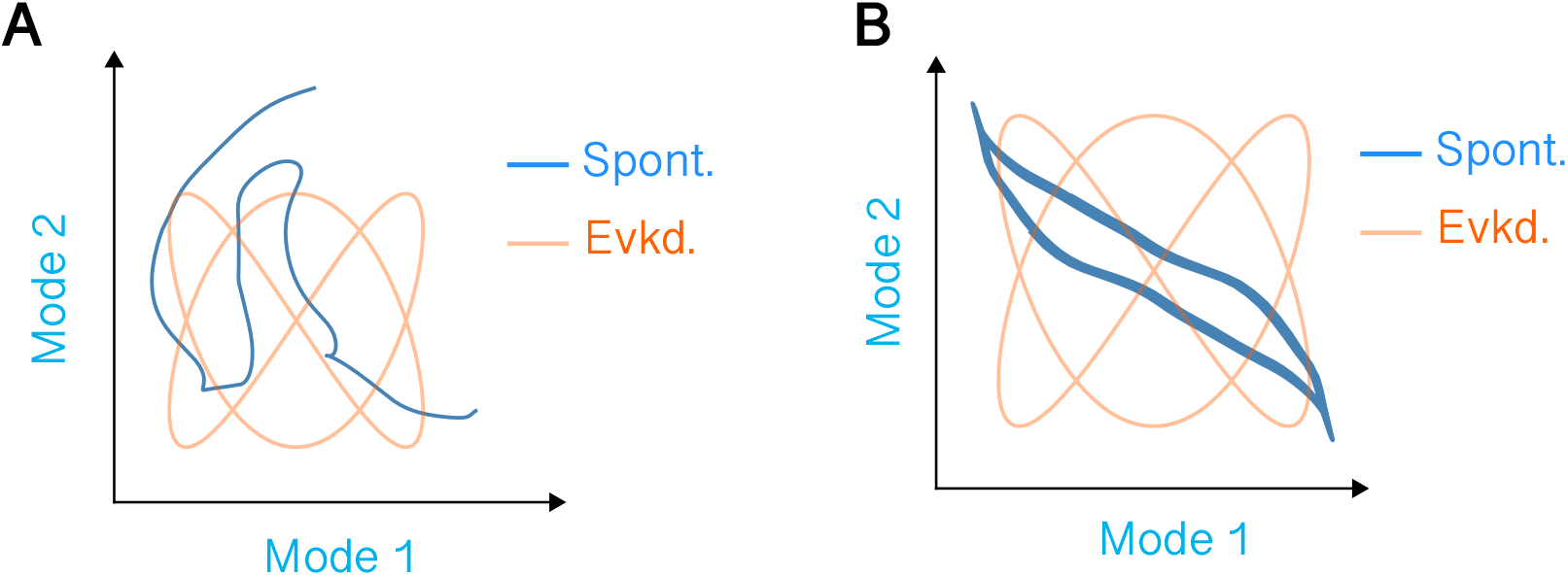
Effect of temporal filter ablations on spontaneous network dynamics. Low-dimensional population dynamics of evoked activity (orange) and spontaneous activity (blue). (A) Removal of the postsynaptic integration filter resulted in high-dimensional, irregular spontaneous trajectories with no resemblance to the evoked manifold. (B) Removal of the presynaptic low-pass filter preserved low-dimensional structure in spontaneous activity but caused a misalignment with the evoked trajectories.

## References

1. Arieli, A., Sterkin, A., Grinvald, A., & Aertsen, A. (1996). Dynamics of ongoing activity: explanation of the large variability in evoked cortical responses. Science, 273(5283), 1868–1871.

2. Asabuki, T., & Clopath, C. (2025). Taming the chaos gently: a predictive alignment learning rule in recurrent neural networks. Nature Communications, 16(1), 6784.

3. Asabuki, T., & Fukai, T. (2020). Somatodendritic consistency check for temporal feature segmentation. Nature Communications, 11(1), 1554.

4. Avitan, L., Pujic, Z., Mölter, J., Zhu, S., Sun, B., & Goodhill, G. J. (2021). Spontaneous and evoked activity patterns diverge over development. Elife, 10, e61942.

5. Bastos, A. M., Usrey, W. M., Adams, R. A., Mangun, G. R., Fries, P., & Friston, K. J. (2012). Canonical microcircuits for predictive coding. Neuron, 76(4), 695–711.

6. Berkes, P., Orbán, G., Lengyel, M., & Fiser, J. (2011). Spontaneous cortical activity reveals hallmarks of an optimal internal model of the environment. Science, 331(6013), 83–87.

7. Breakspear, M. (2017). Dynamic models of large-scale brain activity. Nature Neuroscience, 20(3), 340–352.

8. Chalk, M., Marre, O., & Tkačik, G. (2018). Toward a unified theory of efficient, predictive, and sparse coding. Proceedings of the National Academy of Sciences (USA), 115(1), E859–E868.

9. Churchland, M. M., Cunningham, J. P., Kaufman, M. T., Foster, J. D., Nuyujukian, P., Ryu, S. I., & Shenoy, K. V. (2012). Neural population dynamics during reaching. Nature, 487(7405), 51–56.

10. Cunningham, J. P., & Yu, B. M. (2014). Dimensionality reduction for large-scale neural recordings. Nature neuroscience, 17(11), 1500–1509.

11. Deco, G., Jirsa, V. K., & McIntosh, A. R. (2013). Emerging concepts for the dynamical organization of resting-state activity in the brain. Nature Reviews Neuroscience, 14(5), 336–349.

12. Dimakou, A., Pezzulo, G., Zangrossi, A., & Corbetta, M. (2025). The predictive nature of spontaneous brain activity across scales and species. Neuron, 113(9), 1310–1332.

13. Elsayed, G. F., Lara, A. H., Kaufman, M. T., Churchland, M. M., & Cunningham, J. P. (2016). Reorganization between preparatory and movement population responses in motor cortex. Nature communications, 7(1), 13239.

14. Fiser, J., Berkes, P., Orbán, G., & Lengyel, M. (2010). Statistically optimal perception and learning: from behavior to neural representations. Trends in cognitive sciences, 14(3), 119–130.

15. Fiser, J., Chiu, C., & Weliky, M. (2004). Small modulation of ongoing cortical dynamics by sensory input during natural vision. Science, 303(5664), 967–972.

16. Foster, D. J., & Wilson, M. A. (2006). Reverse replay of behavioural sequences in hippocampal place cells during the awake state. Nature, 440(7084), 680–683.

17. Fox, M. D., & Raichle, M. E. (2007). Spontaneous fluctuations in brain activity observed with functional magnetic resonance imaging. Nature reviews neuroscience, 8(9), 700–711.

18. Friston, K. (2010). The free-energy principle: a unified brain theory? Nature Reviews Neuroscience, 11(2), 127–138.

19. Gallego, J. A., Perich, M. G., Miller, L. E., & Solla, S. A. (2017). Neural manifolds for the control of movement. Neuron, 94(5), 978–984.

20. Huang, Y., & Rao, R. P. N. (2011). Predictive coding. Neuron, 72(4), 557–576.

21. Iyer, K. K., Hwang, K., Hearne, L. J., Muller, E., D’Esposito, M., Shine, J. M., & Cocchi, L. (2022). Focal neural perturbations reshape low-dimensional trajectories of brain activity supporting cognitive performance. Nature communications, 13(1), 4.

22. Kenet, T., Bibitchkov, D., Tsodyks, M., Grinvald, A., & Arieli, A. (2003). Spontaneously emerging cortical representations of visual attributes. Nature, 425(6961), 954–956.

23. Koren, V., & Denève, S. (2017). Computational account of spontaneous activity as a signature of predictive coding. PLoS Computational Biology, 13(1), e1005355.

24. Levenstein, D., Efremov, A., Eyono, R. H., Peyrache, A., & Richards, B. (2024). Sequential predictive learning is a unifying theory for hippocampal representation and replay. BioRxiv, 2024–04.

25. Litwin-Kumar, A., & Doiron, B. (2012). Slow dynamics and high variability in balanced cortical networks with clustered connections. Nature neuroscience, 15(11), 1498–1505.

26. Luczak, A., Bartho, P., & Harris, K. D. (2009). Spontaneous events outline the realm of possible sensory responses in neocortical populations. Neuron, 62(3), 413–425.

27. Luczak, A., Bartho, P., & Harris, K. D. (2013). Gating of sensory input by spontaneous cortical activity. Journal of Neuroscience, 33(4), 1684–1695.

28. Luczak, A., McNaughton, B. L., & Kubo, Y. (2022). Neurons learn by predicting future activity. Nature Machine Intelligence, 4(1), 62–72.

29. Marguet, S. L., & Harris, K. D. (2011). State-dependent representation of amplitude-modulated noise stimuli in rat auditory cortex. Journal of Neuroscience, 31(17), 6414–6420.

30. Mazzucato, L., Fontanini, A., & La Camera, G. (2019). Stimuli-response variability in cortical networks with multiple stable states. Current Opinion in Neurobiology, 58, 60–69.

31. Orbán, G., Berkes, P., Fiser, J., & Lengyel, M. (2016). Neural variability and sampling-based probabilistic representations in the visual cortex. Nature Neuroscience, 19(1), 58–64.

32. Pezzulo, G., Zorzi, M., & Corbetta, M. (2021). The secret life of predictive brains: what’s spontaneous activity for? Trends in Neurosciences, 44(9), 775–788.

33. Rabinovich, M. I., Huerta, R., & Laurent, G. (2008). Transient dynamics for neural processing. Science, 321(5885), 48–50.

34. Raichle, M. E. (2006). The brain’s dark energy. Science, 314(5803), 1249–1250.

35. Rao, R. P., & Ballard, D. H. (1999). Predictive coding in the visual cortex: a functional interpretation of some extra-classical receptive-field effects. Nature neuroscience, 2(1), 79–87.

36. Recanatesi, S., Farrell, M., Lajoie, G. et al. Predictive learning as a network mechanism for extracting low-dimensional latent space representations. Nat Commun 12, 1417 (2021).

37. Ringach, D. L. (2009). Spontaneous and driven cortical activity: implications for computation. Current opinion in neurobiology, 19(4), 439–444.

38. Runfola, C., Petkoski, S., Sheheitli, H., Bernard, C., McIntosh, A. R., & Jirsa, V. (2025). A mechanism for the emergence of low-dimensional structures in brain dynamics. npj Systems Biology and Applications, 11(1), 32.

39. Sadtler, P. T., Quick, K. M., Golub, M. D., Chase, S. M., Ryu, S. I., Tyler-Kabara, E. C., … & Batista, A. P. (2014). Neural constraints on learning. Nature, 512(7515), 423–426.

40. Saxena, S., & Cunningham, J. P. (2019). Towards the neural population doctrine. Current opinion in neurobiology, 55, 103–111.

41. Sengupta, B., Laughlin, S. B., & Niven, J. E. (2013). Balanced excitatory and inhibitory currents underlie efficient coding and metabolic efficiency. PLoS Computational Biology, 9(10), e1003263.

42. Shenoy, K. V., Sahani, M., & Churchland, M. M. (2013). Cortical control of arm movements: a dynamical systems perspective. Annual review of neuroscience, 36(1), 337–359.

43. Sompolinsky, H., Crisanti, A., & Sommers, H. J. (1988). Chaos in random neural networks. Physical review letters, 61(3), 259.

44. Song, H., Shim, W. M., & Rosenberg, M. D. (2023). Large-scale neural dynamics in a shared low-dimensional state space reflect cognitive and attentional dynamics. Elife, 12, e85487.

45. Stringer, C., Pachitariu, M., Steinmetz, N., Carandini, M., & Harris, K. D. (2019). High-dimensional geometry of population responses in visual cortex. Nature, 571(7765), 361–365.

46. Tsodyks, M., Kenet, T., Grinvald, A., & Arieli, A. (1999). Linking spontaneous activity of single cortical neurons and the underlying functional architecture. Science, 286(5446), 1943–1946.

47. Uddin, L. Q. (2020). Bring the noise: reconceptualizing spontaneous neural activity. Trends in cognitive sciences, 24(9), 734–746.

48. Wilson, M. A., & McNaughton, B. L. (1994). Reactivation of hippocampal ensemble memories during sleep. Science, 265(5172), 676–679.

49. Yu, B. M., Cunningham, J. P., Santhanam, G., Ryu, S., Shenoy, K. V., & Sahani, M. (2008). Gaussian-process factor analysis for low-dimensional single-trial analysis of neural population activity. Advances in neural information processing systems, 21.

50. de Oliveira, E. F., Kim, S., Qiu, T. S., Peyrache, A., Batista-Brito, R., & Sjulson, L. (2022). Off-manifold coding in visual cortex revealed by sleep. Biorxiv, 2022–06.

